# FARESHARE: An Open-source Apparatus for Assessing Drinking Microstructure in Socially Housed Rats

**DOI:** 10.1101/2023.10.06.561248

**Authors:** Jude A. Frie, Jibran Y. Khokhar

## Abstract

Social factors have been shown to play a significant and lasting role in alcohol consumption. Studying the role of social context on alcohol drinking is important to understand the factors that contribute to initiation or maintenance of casual and problematic alcohol use, as well as those that may be protective. A substantial body of preclinical research has shown that social environment such as housing conditions and social rank plays an important role in alcohol consumption and preference, though the extent of these effects have been obfuscated by methodological differences and technical challenges. Robust individual differences in alcohol intake in socially housed animals are difficult to track when animals share a common fluid source. Commercial solutions are prohibitively expensive and are limited by proprietary software and hardware (including caging systems). Here we describe an affordable, open-source solution for tracking fluid consumption in socially housed rats.

## Introduction

Given the lack of new effective therapeutics in addiction treatment despite a substantial and rich mechanistic preclinical literature, there has been a recent effort in to improve the translational validity of animal models by increasing the ecological relevance of experiments (Venniro et al., 2020). Previous findings have shown that risk of developing substance use, and the effectiveness of treatment interventions depend greatly on social environment (Knight and Simpson, 1996). The role of social factors such as peer pressure, loneliness, and social stress and support on alcohol consumption and related problems has been a large focus in clinical alcohol research, with significant facilitation of binge drinking by peers, increased risk of alcohol problems in solitary drinkers, and gender differences in motives for alcohol consumption in social settings (Freisthler et al., 2014; Skrzynski and Creswell, 2021; Smit et al., 2015). Thus, incorporating social context into preclinical models of substance use is an important step toward more translational findings.

Preclinical studies have shown that the presence of other animals may facilitate or reduce alcohol drinking depending on the particular social environment such as housing conditions and social rank (PMID: 20616986); however these effects are inconsistent due to methodological inconsistencies and technical challenges. It is difficult to track individual differences in consumption when animals share a cage and common substance source. The majority of drug intake studies in rodents have required social isolation of subjects to get around this issue, neglecting the social context so relevant to clinical populations (Knight and Simpson, 1996; Renda et al., 2020). Previous attempts to track fluid consumption in socially housed animals can be placed into the following categories: 1) Track total consumption of the group (Ehlers et al., 2007; Juárez and Vázquez-Cortés, 2003; Wolffgramm and Heyne, 1991), 2)use a mesh cage separator (Al-Sabagh et al., 2022; Hamidullah et al., 2021; Palm and Nylander, 2014; Rizk et al., 2022; Tomie et al., 2007), 3) video recording (Logue et al., 2014; Varlinskaya et al., 2015), or 4) radio frequency tracking (Terem et al., 2022; Walcott and Ryabinin, 2021; Woodard et al., 2020). Tracking total consumption is the easiest method but 2) loses all temporal resolution and cannot evaluate individual differences or effect of social rank (Ryabinin and Walcott, 2018). Mesh cage separators similarly have no temporal resolution and are difficult to interpret as they do not allow for naturalistic social interaction. Video tracking software can measure the time rats spend close to the sipper, but whether this time is associated with drinking is not clear. Video tracking also struggles to distinguish between animals that are close in proximity to each other or maintaining identity after close contacts. Radio frequency tracking is the most promising solution and has been used in commercial products such as IntelliCage. IntelliCage, however, is prohibitively expensive (∼$65,000 USD per cage) and as a commercial product is limited by proprietary software and hardware, including the incompatibility with existing caging systems. Additionally, individual drinking is measured with only a lickometer, and while licking has been shown to be correlational with volume of consumption, the number of licks in any given bout may vary with drinking context (Ryabinin and Walcott, 2018).

It is clear that there is a need for a non-commercial, open-source alternative to improve the quality and availability of preclinical alcohol research in social settings. One such solution is SIP, a project that continuously monitors polysubstance fluid intake in group housed mice (Wong et al., 2023). This project benefits from delivery of fluid when mice are present rather than passive sipping, thus exact volume can be measured and drinking microstructure may be recorded and analysed. Unfortunately, without a lickometer, it cannot be certain that the animal is actually drinking the delivered volume. Additionally, SIP is specialized toward polysubstance intake in mice, still uses expensive, proprietary fluid delivery systems and radio-frequency identification (RFID) hardware and software and requires a custom-built home cage. Another project, PiDose, uses RFID, lickometer, and fluid delivery thus ensuring that volumes delivered are consumed, and accurate volume information is collected (Woodard et al., 2020). However, fluid intake is not truly social as PiDose has a separate chamber for drug delivery. PiDose is also specialized for giving specific doses of drug to animals based on weight and uses different delivery methods for drug and water. RFID has also recently been implemented with a Bpod state machine to successfully monitor self-administration of fentanyl paired with fibre photometry in group housed mice (Terem et al., 2022), but also uses a tunnel-based design with only one mouse being able to access both fluids at one time. These projects are important contributions to preclinical research, though a general use, low-cost method for measuring individual momentary fluid intake in socially housed rats is warranted. Thus, we developed a device for **F**luid **A**cquisition **RE**cording in **S**ocially **H**oused **A**nimal **RE**search (FARESHARE) to allow for RFID-based fluid tracking in rats.

FARESHARE is an affordable, open-source device that can be placed in any home cage or operant box that can be used with as many animals as required for the study design. It uses short-range RFID to identify individual rats, a lickometer to determine when animals are drinking to begin fluid delivery, custom low-profile PCB that sits directly on top of an Arduino-based microcontroller, volumetric fluid delivery via a custom peristaltic pump for accurate measurement of consumption volume (that may also be used for other purposes), OLED display for showing overall individual fluid consumption and licks, and continuous data logging to an SD card module. Here we validate FARESHARE via a two-bottle choice paradigm using two FARESHARE devices: one for water, and one for alcohol. We use the data collected to determine fluid consumption and drinking microstructure for each of four rats housed in the same cage. Having a robust, affordable method for measuring drinking microstructure in socially housed animals will be of considerable benefit in preclinical addiction research, and a step toward more translationally and ecologically relevant animal models of fluid consumption. The added dimension of time allows for the analysis of circadian-linked consumption and the discrimination of binge-like drinking behaviours. Additionally, the open-source nature of the project enables researchers to customize the device for more advanced applications such as sending signals to additional peripherals (e.g., optogenetic stimulation or electrophysiology/fiber photometry) or software on drinking initiation for time-locked or closed-loop interventions, manipulations, and measurements.

## Materials and Methods

### Animals

Four female Sprague Dawley rats (Charles River Lab, St. Constant, Canada) were kept on a 12:12 light-dark cycle (lights off at 0800 hours) in a colony maintained at 21°C in a single polyethylene Guinea pig cage. Food (Envigo, Madison, Wisconsin, USA, Rodent Diet, 14% protein) and water were available ad libitum. All animal procedures were performed in accordance with the University of Guelph animal care committee, the University of Western Ontario Animal Use Subcommittee, and were consistent with guidelines established by the Canadian Council on Animal Care.

### Injection of RFID Tags

Rats were injected subcutaneously with 5mg/kg carprofen prior to RFID injection. Anesthesia was induced with 4-5% isoflurane. RFID tags were injected using an RFID chip injector under the scalp and RFID was placed such that the tip of the tag sat between the eyes and extended back toward the ears as shown in figure 7.

### Design of Electronics and 3D Printer Files

FARESHARE uses the Arduino-based RedBoard Qwiic that has a ATmega328 processor with a 16MHz Clock Speed, 20 I/O pins, and 32kb of flash memory. All code was written in Arduino language using Arduino IDE with added open-source libraries: Adafruit GFX Library, Adafruit SSD1306, Sparkfun Qwiic OpenLog, and CapacitiveSensor. The PCB was designed in the open-source CAD software Fritzing. The PCB acts as the bridge between the RedBoard, RFID reader (ID-12LA), OLED display, SD card reader, Capacitive lickometer, and motor controller. An A4988 Stepper Motor Driver controls the peristaltic pump’s Nema 17 Stepper Motor. 3D printer files were created in SolidWorks and printed on a Creality K1 Speedy printer using PLA filament with Creality Print slicer.

### Pump Characterization

To evaluate the error between volume reading and actual volume, the output tube of one device was placed in a beaker and the beaker was placed on an analytical scale. The device was then activated until 1.00mL was output on the OLED display and the weight reading on the scale was recorded. The scale was then tared, and another mL was measured. This was repeated 20 times without resetting the FARESHARE to determine if error would be stable over time.

### Measurement of Alcohol consumption and Preference

Two reservoirs were prepared in 1L Erlenmeyer flasks (10% ethanol v/v and tap water). Each flask was sealed with parafilm except for a small hole for the FARESHARE input pump hose to go through. Two FARESHARE devices were attached to the inside of one wall of the home cage and primed for either alcohol or water. Rats were then given 24-hour access to both devices for nine days.

## Data Accessibility

All Arduino code, GERBER files, PCB design file, and STLs for 3D printed parts are available at https://github.com/jfrie/FARESHARE.

### Statistical Analysis

Statistics were conducted in GraphPad Prism 9.5.1. Pump error was evaluated by simple linear regression and descriptive statistics. Changes in daily alcohol consumption and preference were analyzed via one-way ANOVA with day as a repeated measure. Individual hourly water and alcohol consumption were evaluated via two-way ANOVA with time as a repeated measure. Effect of light cycle and bout range on each subject’s water and alcohol consumption was evaluated via two-way ANOVA. Max and mean bout sizes, volume per lick, and bout frequency between fluid types were analyzed via two-tailed paired t-test. Bouts were defined as a drinking event containing at least 20 licks, similar to previous literature (Eastwood et al., 2014; Ford et al., 2009; Kampov-Polevoy et al., 2000; Samson et al., 1991). A p-value<0.05 was considered significant for all comparisons. Results are displayed as mean ± SEM.

### Build Instructions

PCB Ordering: Upload GERBER file to https://jlcpcb.com and order with default settings.

**Table 1:** Bill of Materials.

| Part Name | Distributor | Alternate | Cost (\$US) |
| --- | --- | --- | --- |
| 1000uF Capacitor (through hole) | Mouser | Digikey | 1.63 |
| 16G Needles (x2) | Fisher Scientific | Surgo | 0.85 |
| 1M Ohm Resistor (through hole) | Mouser | Digikey | 0.42 |
| 5-pin 2.54mm JST Female to Female Cable | Walmart | Amazon | 1.30 |
| 5-pin 2.54mm JST Male header (x2) | Walmart | Amazon | 0.14 |
| 3D printer filament | Amazon | Hatchbox | 2.50 |
| A4988 Stepper Motor Driver | Amazon | Digikey | 1.20 |
| Ball bearings (3mm x 8mm x 4mm) (x4) | Amazon | NewEgg | 2.00 |
| Barrel Jack PCB mount (3 pin) | Mouser | Digikey | 1.25 |
| Break Away Headers (2.54mm straight) (41 pins) | Sparkfun | Mouser | 3.50 |
| Custom PCB | JLCPCB | OSHPark | 2.00 |
| Dual lock (10cm) | Amazon | Mouser | 1.50 |
| Epoxy | Walmart | Amazon | 6.12 |
| Hook-up Wire (22 AWG) | Sparkfun | Amazon | 2.95 |
| M3 Screw (Hex Socket, 25mm) (x4) | Home Hardware | Amazon | 2.75 |
| M3 Screw (Hex Socket, 10mm) (x7) | Home Hardware | Amazon | 3.25 |
| M3 Screw (Hex Socket, 5mm) (x8) | Home Hardware | Amazon | 3.50 |
| M3 Nut (x3) | Home Hardware | Amazon | 2.00 |
| Masterflex® Microbore Pump Tubing, Platinum-Cured Silicone, 1.42mm ID, 3.12mm OD (16cm long) | Avantor | Fisher Scientific | 2.30 |
| Mini 4-pin Push Button | Mouser | Amazon | 1.00 |
| Micro SD Card (FAT16 or 32, 64MB to 32GB) | Amazon | Walmart | 5.99 |
| Nema 17 Stepper Motor (4 lead, 2A) (17HS19-2004S1) | Amazon | AliExpress | 13.99 |
| Power Supply (12V, 3A) | Amazon | Walmart | 8.99 |
| Qwiic Cable Kit | Sparkfun | Mouser | 8.95 |
| Qwiic OLED Display (0.91 in, 128x32) | Sparkfun | Mouser | 10.95 |
| Qwiic OpenLog | Sparkfun | Mouser | 18.50 |
| RedBoard Qwiic | Sparkfun | Mouser | 21.50 |
| RFID Chip Injector (2.12x12mm) | Amazon | AliExpress | 6.99 |
| RFID Glass Capsule (125kHz) | Sparkfun | Mouser | 5.50 |
| RFID Reader Breakout | Sparkfun | Mouser | 2.10 |
| RFID Reader ID-12LA (125 kHz) | Sparkfun | Mouser | 32.50 |
| Silicone Tubing 1.5mm ID x 3mm (140 cm) | Amazon | NewEgg | 6.99 |
| Stainless Steel Straw (straight 5mm OD) | Amazon | Whisk | 0.60 |
| **Total** | | | **\$185.71** |

PCB Soldering: The figure below shows the PCB before (left) and after (right) soldering. The PCB is silk screened and labeled such that soldering orientation and positioning is clear.

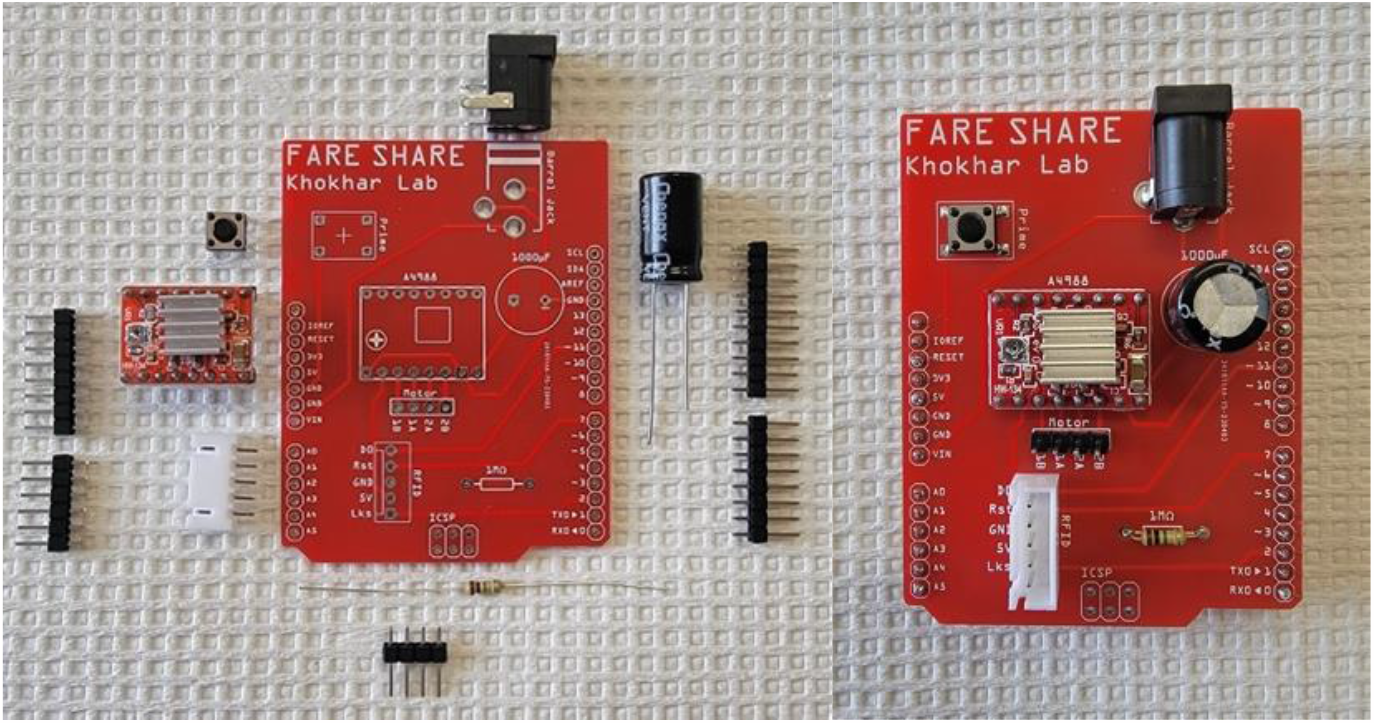

3D Printing: Print in the orientations shown below. The pump rotor and RFID housing should be printed with supports. All other components can be printed without supports. Print with standard PLA filament.

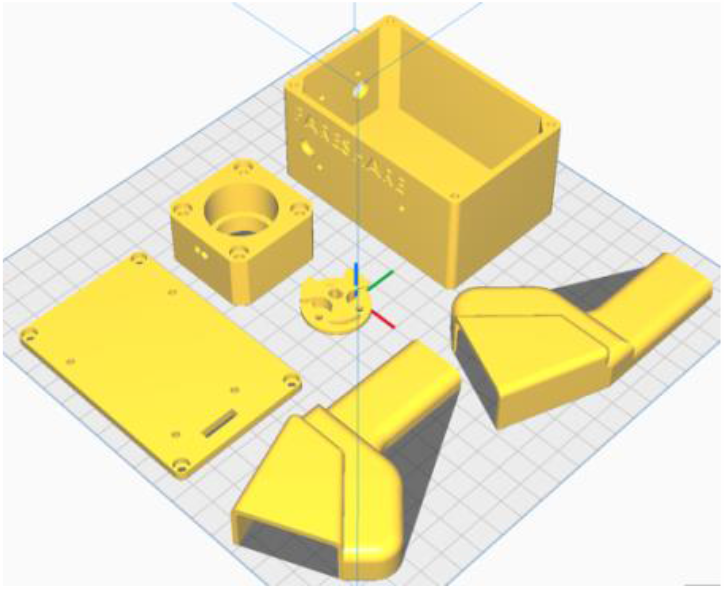

Pump Assembly:

1. Cut two 1cm pieces of needle by slowly rotating and cutting evenly so as not to crimp.

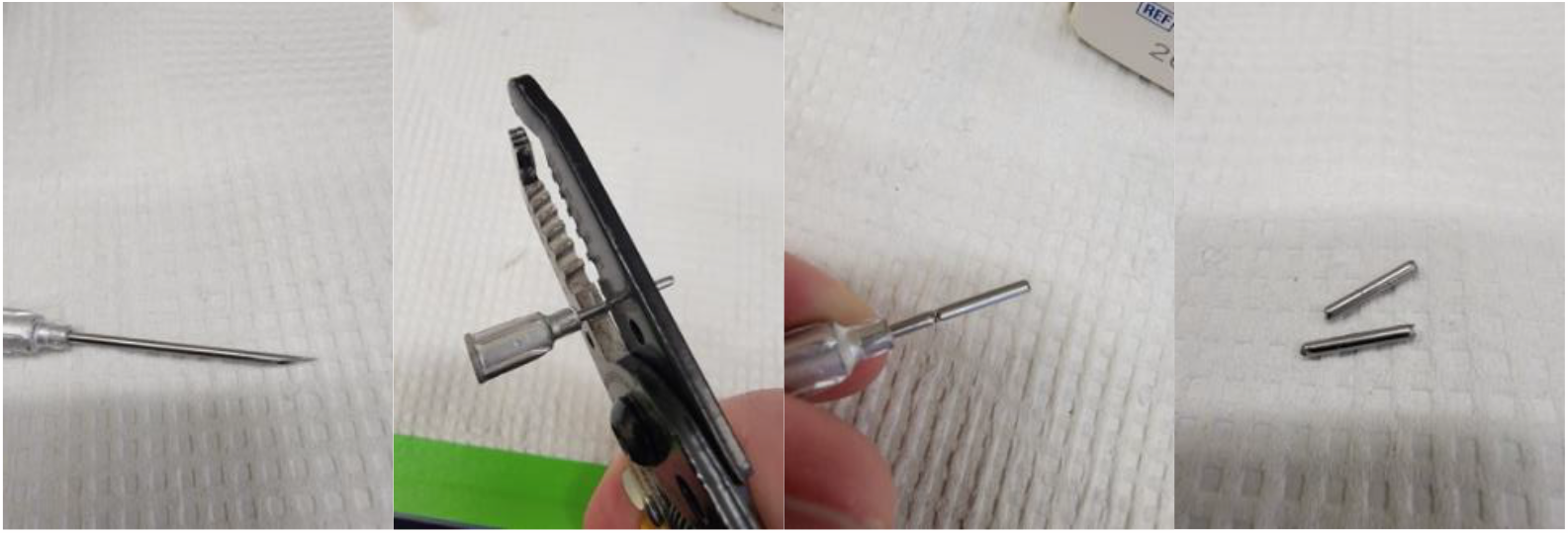
2. Screw M3X10 screws through the holes of the rotor

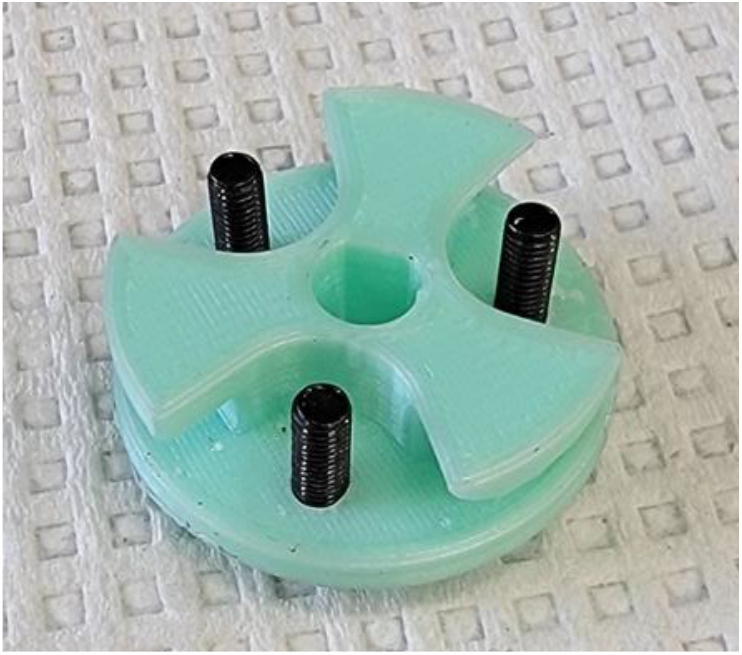
3. Place bearings on screws and secure with M3 nuts. Ensure the nuts are not too tight so as to allow the bearings to turn without any friction.

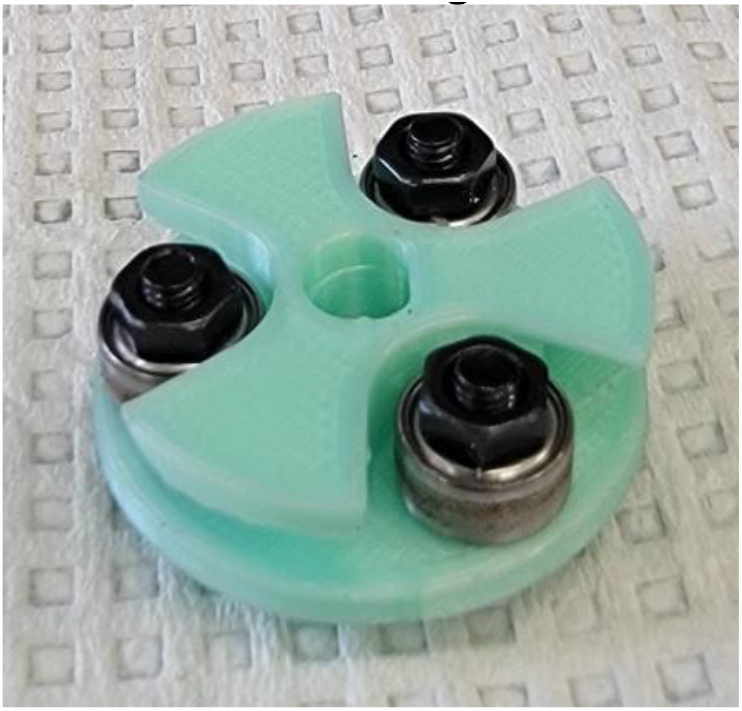
4. Thread 15cm segment of Masterflex tubing through pump case.

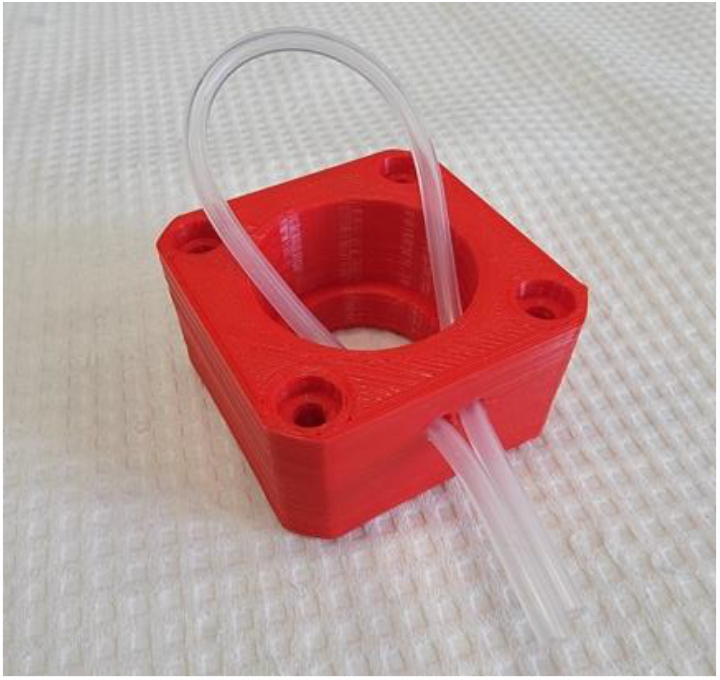
5. Wrap the tubing around the rotor and push the rotor into the pump housing while keeping tension on the tubing. Once inside the housing, manually rotate the rotor a bit and pull on each side of the tubing until the rotor is straight and the tubing wraps around each bearing.

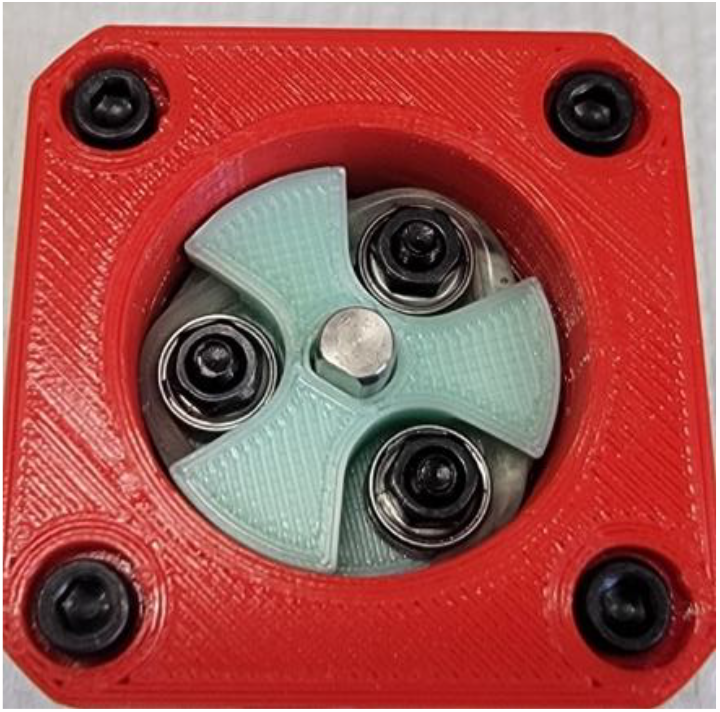
6. Thread another bearing onto the left (with tubing facing you) exiting tube.

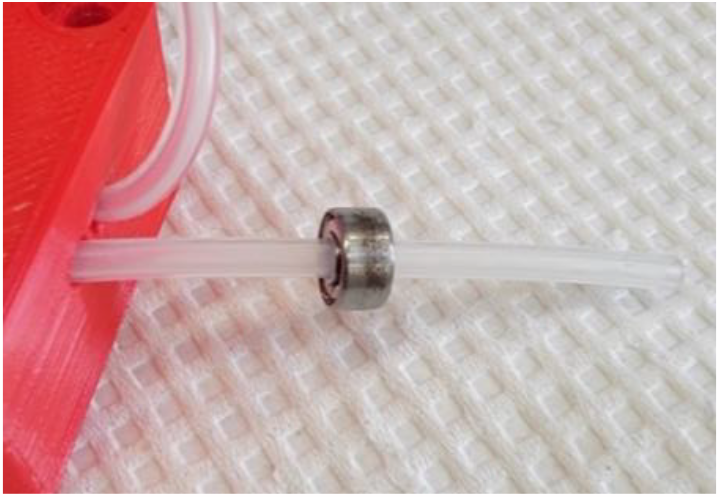
7. Bring bearing up to pump housing and epoxy or super glue to both tubing and housing. This will keep the tubing from being pulled through the rotor or shifting.

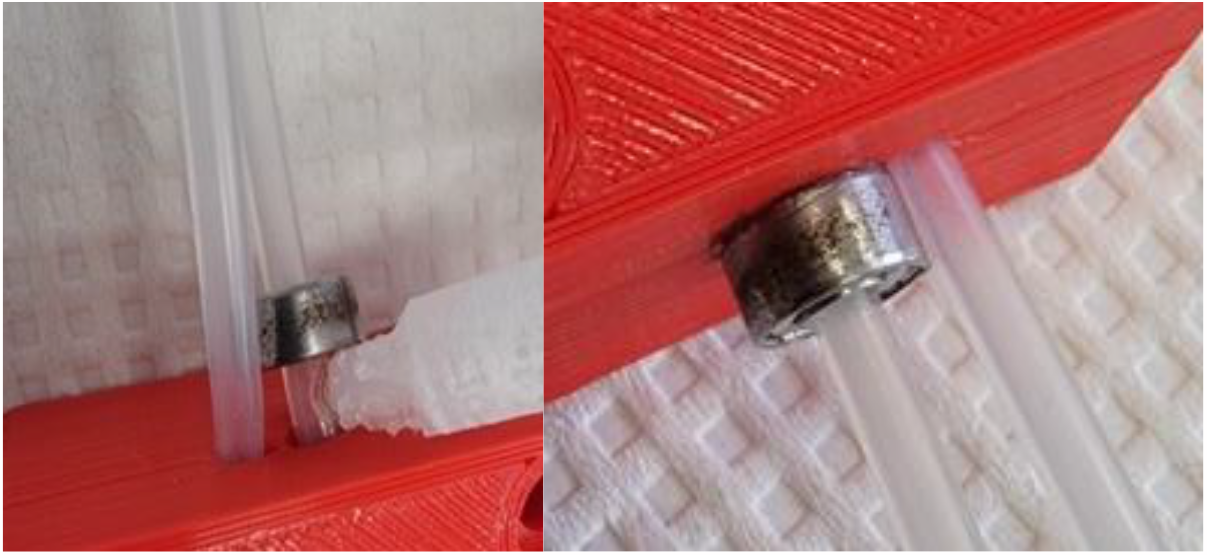
8. Insert needle segments halfway into tubing ends.

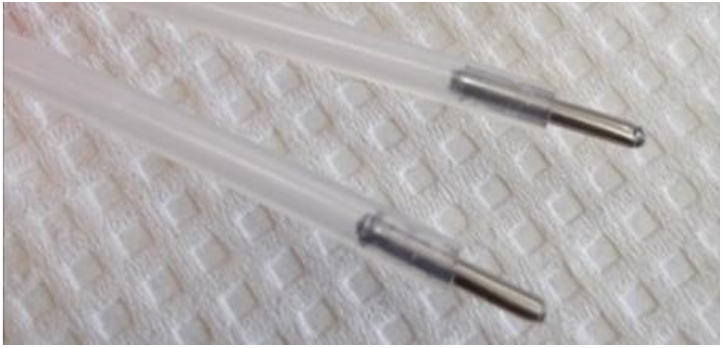
9. Place super glue where needle exits tubing and insert needle ends into two 70 cm segments of silicon tubing (cheaper tubing), gluing tubes together.

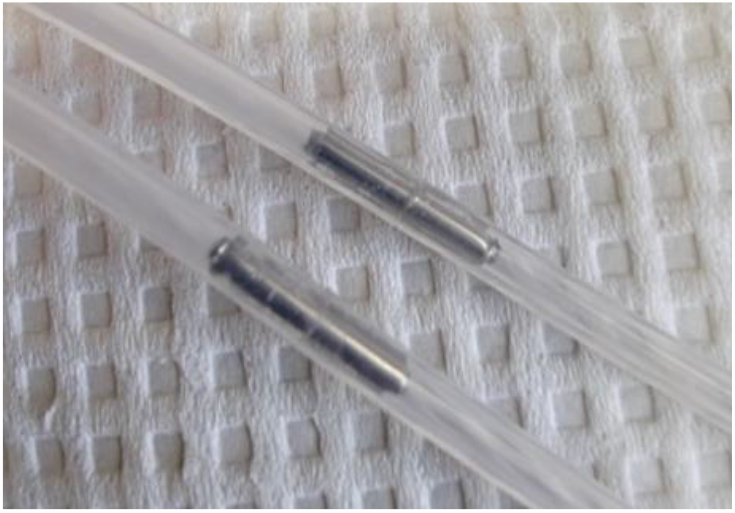
10. Place pump assembly onto motor with tubing facing the same direction as motor wires and, using M3x25mm screws, fix in place.

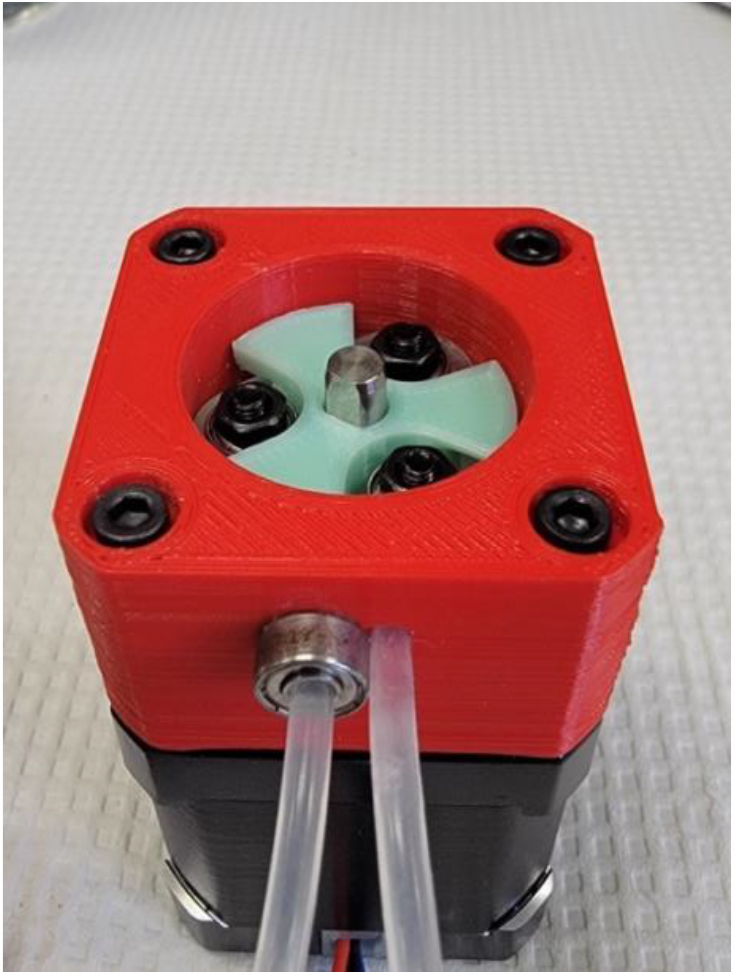
11. Place assembly into box, feeding tubing through input and output holes, and wrapping most of the motor wiring together, placing it in the space between the motor and the wall of the box. Spray the bearings and inner tubing generously with an all-purpose lubricant.

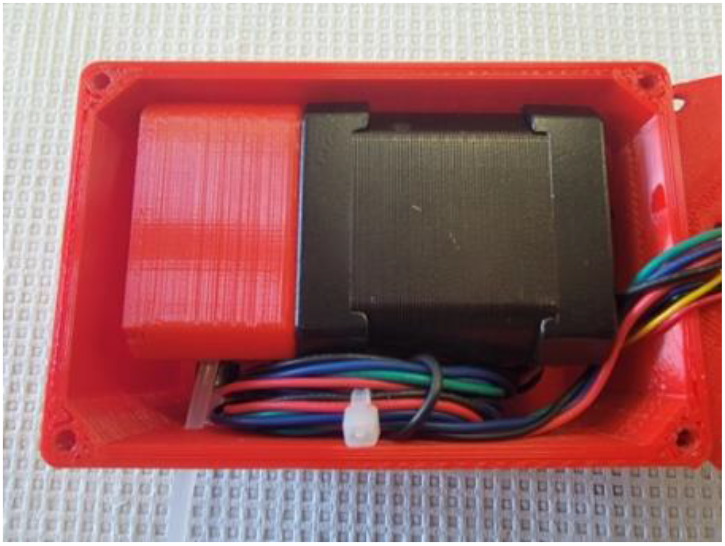
12. Attach OLED display to side of box with M3x4mm screws and thread 10cm Qwiic cable from the left port on the OLED into box.

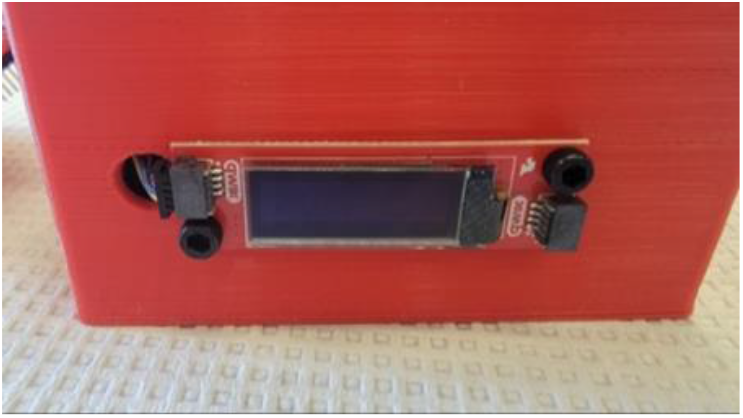
13. Attach data logger to the side of the box with M3x4mm screws and connect Qwiic cable from OLED into right port of data logger. Connect left port to 5cm Qwiic cable.

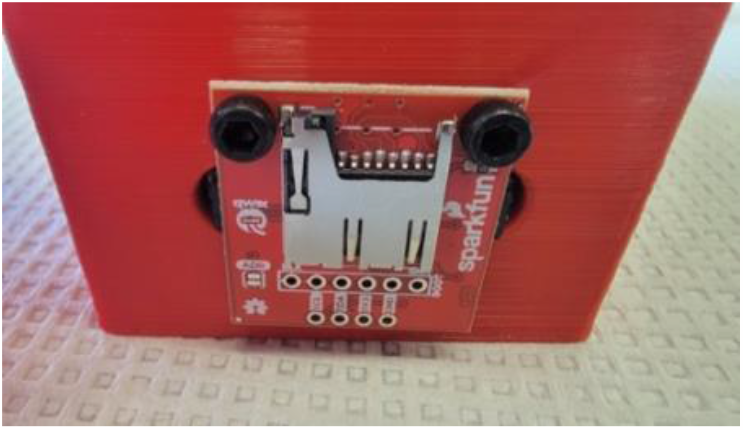
14. Attach RedBoard to box lid with M3x4mm screws. Feed motor wiring and Qwiic cable from data logger through square channel in lid. Connect Qwiic cable to RedBoard. Connect lid to box with M3x10mm screws.

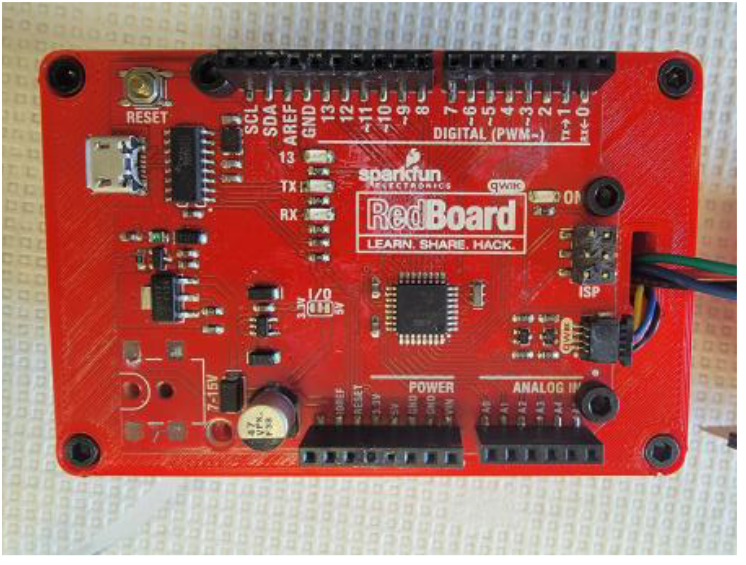
15. Plug PCB into RedBoard and connect motor wires to PCB motor pins with the black wire closest to the resistor.

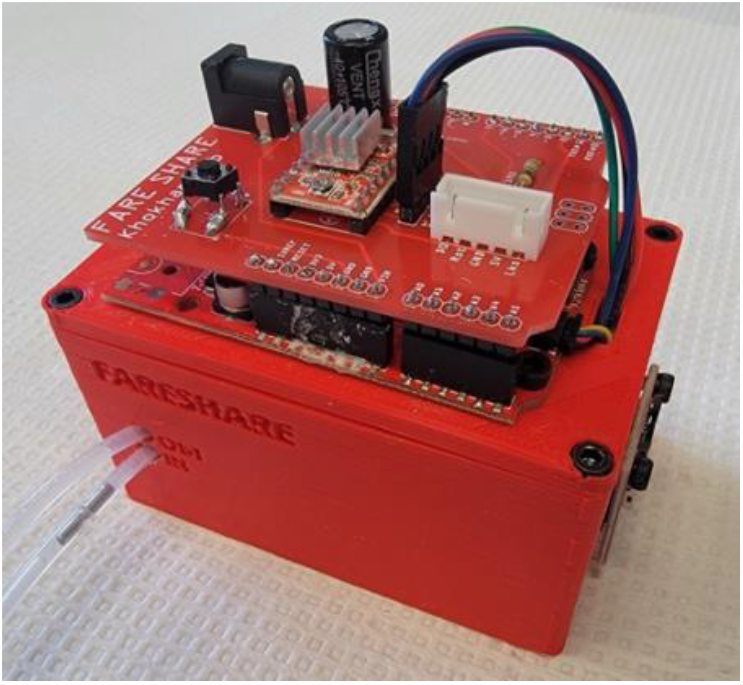

RFID Housing Assembly:

1. Solder RFID Breakout Board to RFID Reader.

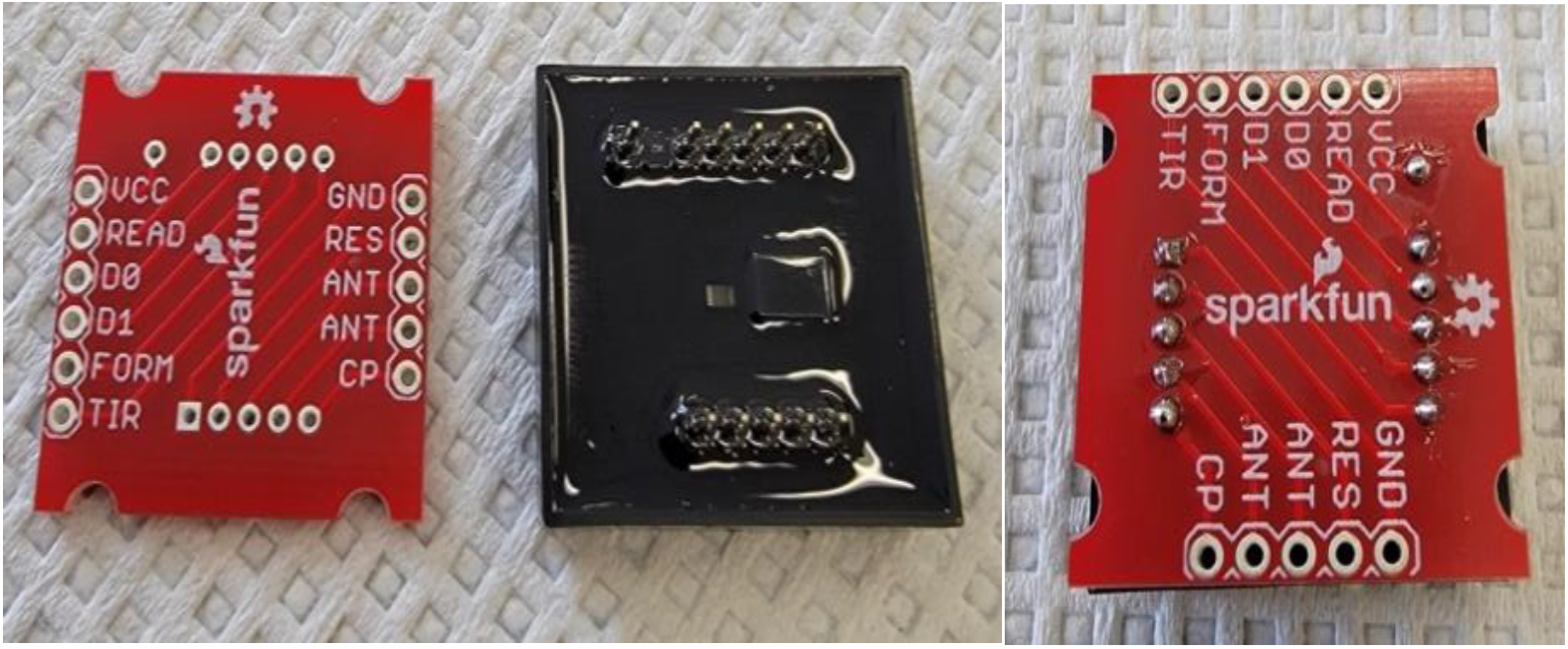
2. Solder a wire between the FORM and GND pins.

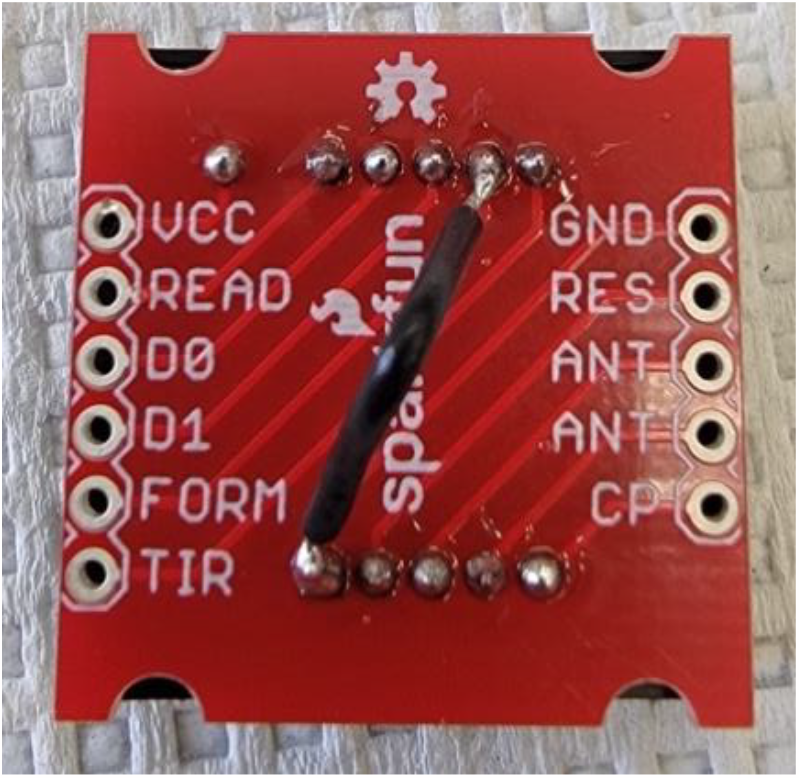
3. Solder 30cm segments of wire to Vcc, D0, GND, and RST through hole pins. Label these wires at the unattached ends so you can identify them later on once they are encased.

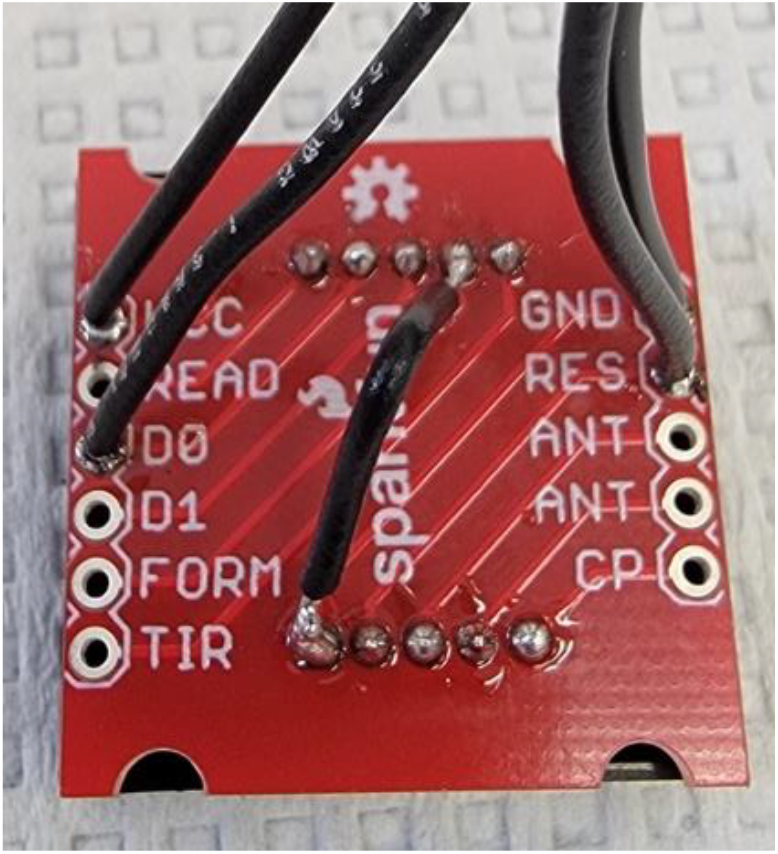
4. Epoxy the metal straw into one half of the RFID housing with 0.5cm of straw projecting past the straw channel. Using glue or command strips, secure RFID sensor inside housing and thread wires through wire channel. Epoxy the other half of RFID housing together.

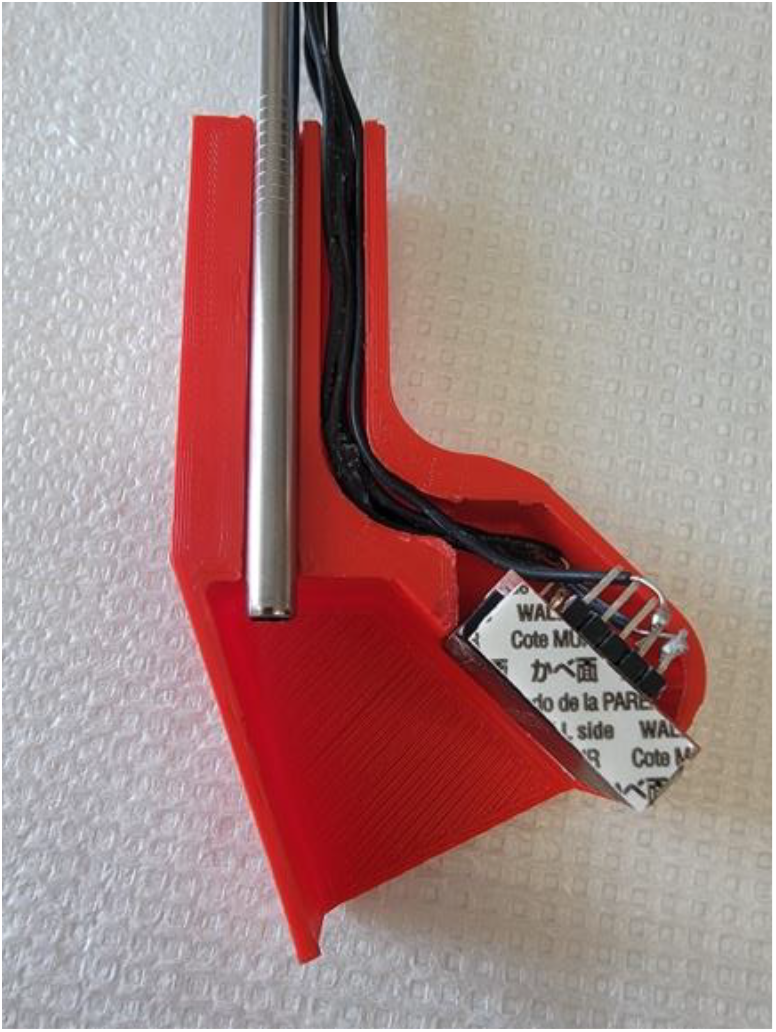
5. Cut the wires such that they are 2cm longer than the straw and, with the same orientation as the plugin on the PCB, solder each wire to a JST header.

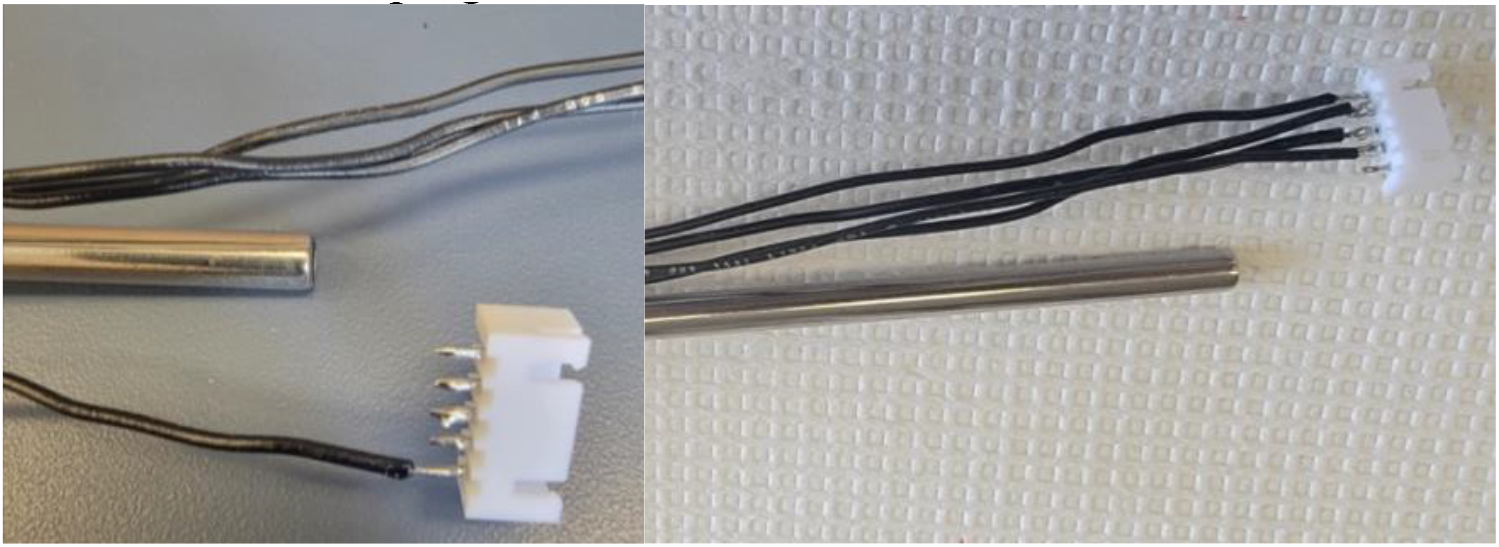
6. Strip 15cm off a 20cm wire. Wrap the stripped section around the tip of the straw and secure with electrical tape. Cut the wire to line up with the remaining pin on the JST header and solder.

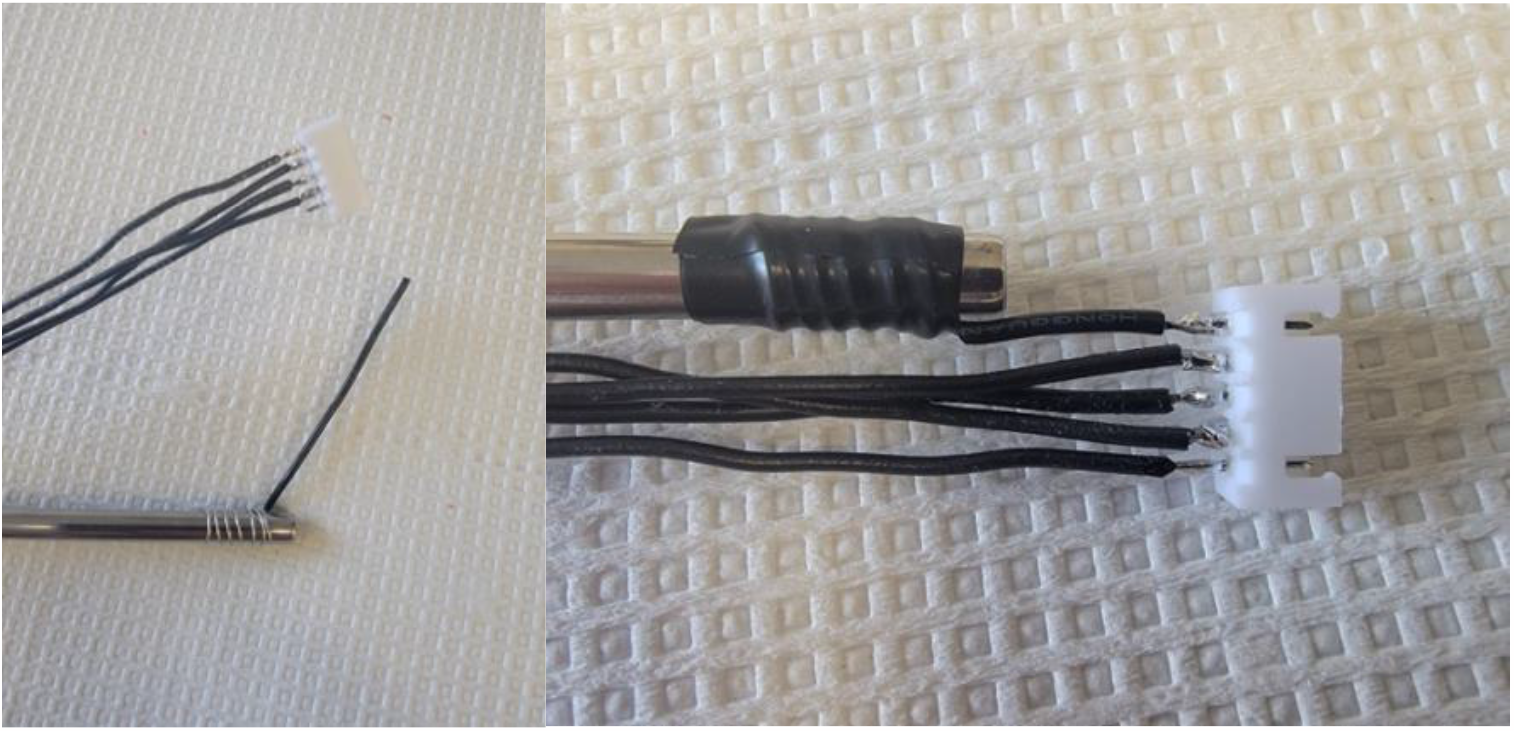
7. Use electrical tape to secure all wire to the straw.

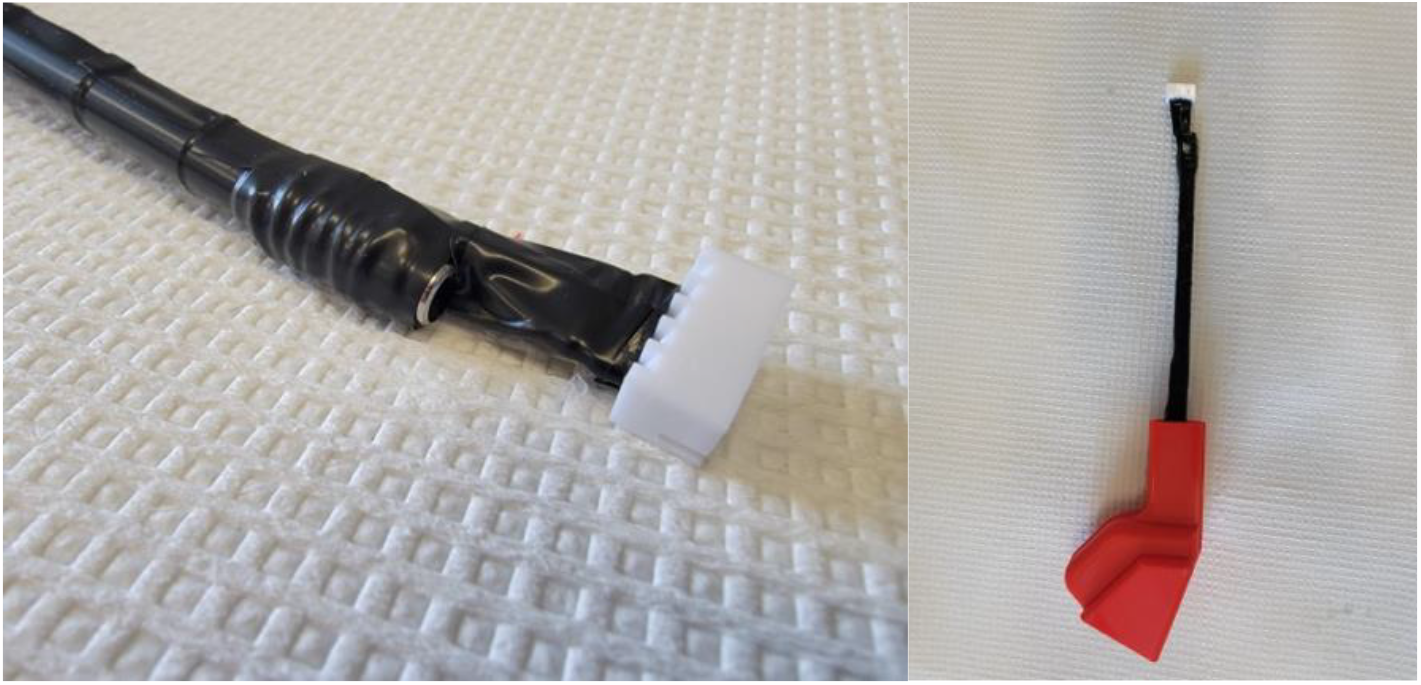

Load libraries: In the Arduino IDE select Tools>Manage Libraries. In the library manager, download Adafruit GFX Library, Adafruit SSD1306, SparkFun Qwiic OpenLog, and CapacitiveSensor.

Addition of RFID Tags to Code: Upload FARESHARE.ino to RedBoard Qwiic. Open serial monitor and place new RFID tag under RFID sensor. The serial monitor will display the new tag ID. Copy this ID and use it to replace the ID on line 38 of the code. Do this for each tag in the following lines for as many rats as are being tracked. Add one additional tag that will be used for priming the pump tubing with fluid when starting an experiment. **Note:** FARESHARE can track as many animals as needed; however, the OLED display only has room to show the overall results of the first four rats.

Pump Calibration: Each assembled peristaltic pump will be slightly different due to small inconsistencies in 3D printing and assembly. Therefore, to ensure each pump delivers an accurate volume and at the same rate, the pumps must be calibrated by the following instructions:

1. Place the input tube into a container of water and the output into a beaker that is at least 10mL.
2. Prime the line using the push button.
3. Place the beaker with the output tube onto an analytical scale.
4. Place an RFID tag by the RFID sensor and tap the straw with your finger to activate the motor. Time how long it takes for FARESHARE to deliver 10g of water.
5. Plug this time in seconds into the following formula: 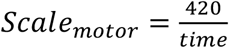 and set the motor_scale variable on line 83 to this new value and upload the new code. This will ensure each device takes the same amount of time to dispense fluid (∼7 minutes for 10mL).
6. Tare the scale and fill it to 10g again with the pump.
7. Plug the volume reading on the FARESHARE OLED display into the following formula: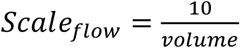 and set the flow_scale variable on line 81 to this new value and upload the new code.

### Operating instructions

1. Prior to starting experiments, remove any .csv or .txt files from the SD card. Insert the SD card into the SD card slot on the Qwiic OpenLog. **Note:** The Qwiic OpenLog SD card logger used in FARESHARE is compatible with 64MB to 32GB microSD cards in either FAT16 or FAT32 formats.
2. Attach straw section of FARESHARE to inside of cage with type 400 dual lock, feeding top of straw through the cage top.
3. Connect wires to PCB via 5-pin JST cable.
4. Place input tube into fluid reservoir and tape in place
5. Plug power supply into the wall and connect the barrel connector to the barrel jack on the PCB. **Note:** Ensure SD card is in reader prior to plugging as files are created when device powers up.
6. Press the reset button on the Arduino to begin the experiment. **Note:** make a note of the time that the experiment is started as the timestamps for each drinking bout are measured in milliseconds following this moment.
7. Place the priming tag onto the RFID sensor to operate the pump until fluid has fully filled the line.
8. Place output tube ∼6cm into straw and tape in place.
9. Allow rats to drink as long as desired.
10. When experiment is complete, unplug FARESHARE and remove SD card.

## Results

### Hardware and Design

FARESHARE requires one standard wall receptacle (110V/60Hz) for power. A single device can be built for ∼$185 US. The device is low profile requiring little lab space to store or use. FARESHARE uses RFID to identify which animal is drinking. When a rat places its head near the RFID scanner and begins licking, an RFID tag implanted under the rat’s scalp will activate a peristaltic pump sending fluid through a metal straw. The straw is capacitive such that it will measure the number of licks in a drinking bout. The total bout volume and lick number are then logged to an SD card. The total volume delivered and total licks since experiment initiation will also be updated in real-time to an OLED display for real-time updates. The SD card creates a .CSV file with timestamped bouts for each rat. The 3D-printed RFID sensor housing is only large enough for a single rat to fit its head, thus, only one rat is ever measured at a time. This design also uniquely allows rats to take turns drinking while in each other’s presence without the need for a door or separate administration chamber to keep the rat isolated for RFID scanning. The system is based around an Arduino Uno type microcontroller to allow labs to easily customize the device for their specific use case. A custom PCB is included to improve circuit reliability, and to make soldering less intensive. All files can be found on GitHub (https://github.com/jfrie/FARESHARE), including 3D printer files, PCB GERBER files, and Arduino code, as well as a bill of materials, detailed build instructions, and operating instructions. A schematic of the device is shown in figure 1.

**Figure 1.**
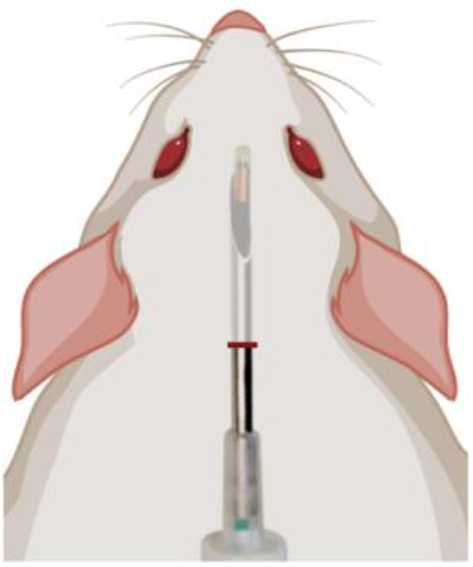
Diagram of RFID positioning and injection sight.

### Pump Characterization

Pump accuracy following 20 measurements of 1.00 mL was high, with an average error of 0.5% (95% CI 0.3-0.7) based on weight. Error also remained stable across tests, with no statistically significant slope following simple linear regression of either raw error (mL; F(1,18)=0.013, P=0.91) or percent error (F(1,18)=2.40, P=0.14) as shown in figure 2. A durability test found the pump was able to run continuously for 30.5 hours before signs of tubing failure. As the cumulative alcohol drinking duration of all rats combined over the nine days was 2.2 hours, this would correspond to a duration of ∼125 days for the current experiment before failure.

**Figure 2.**
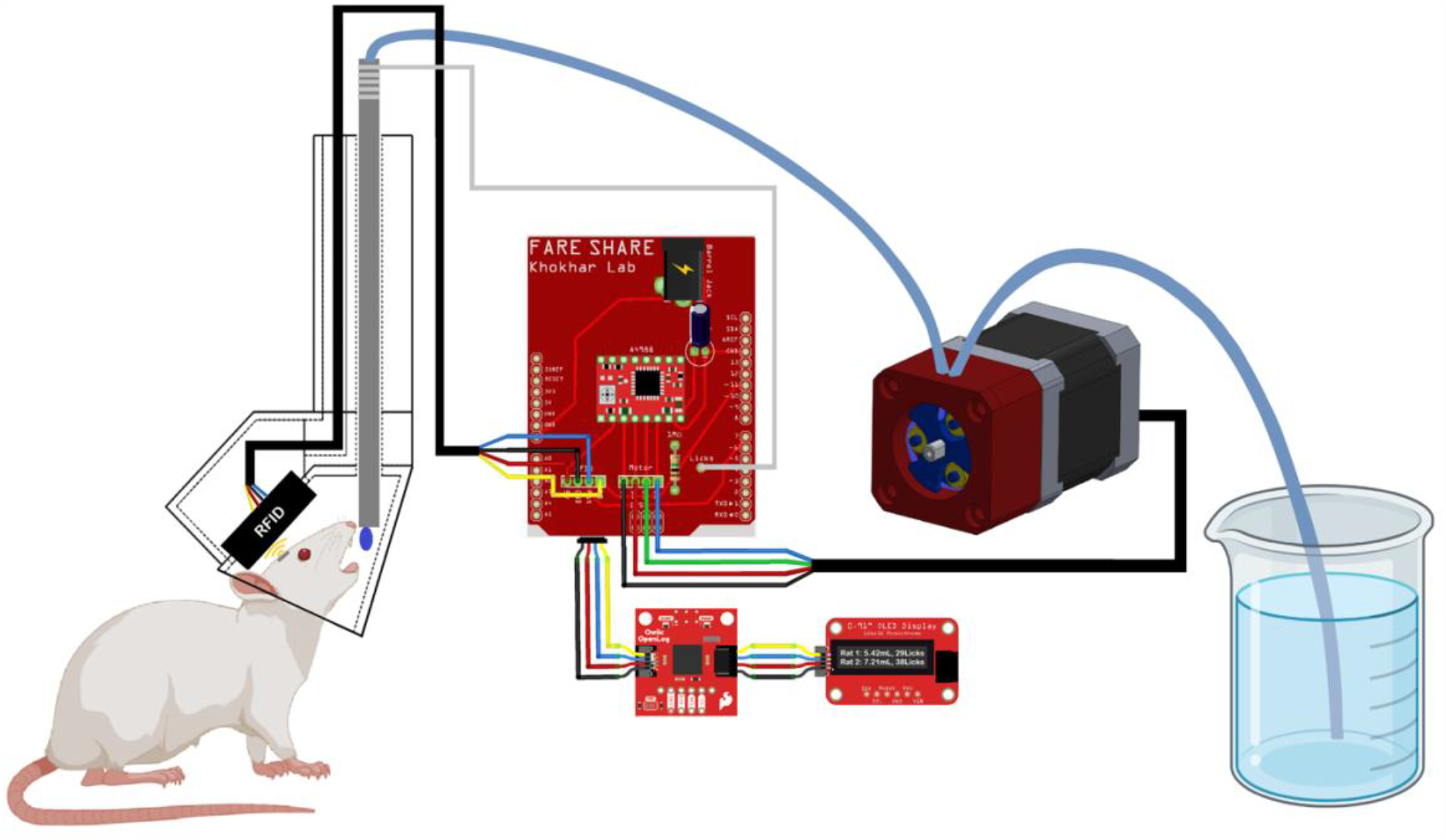
Schematic of FARESHARE. A central Arduino based microcontroller and PCB are connected to a custom peristaltic pump, SD card reader, OLED display, RFID scanner, and capacitive metal straw.

### Fluid Consumption in Group Housed Rats

Rats showed consistent daily alcohol consumption across days as shown in figure 3a. Alcohol preference (figure 3b) did shift across days (effect of day, F(1.854,5.561)=7.455, P=0.028), though no significant differences between days were seen following a Sidak post-hoc test between days.

**Figure 3.**
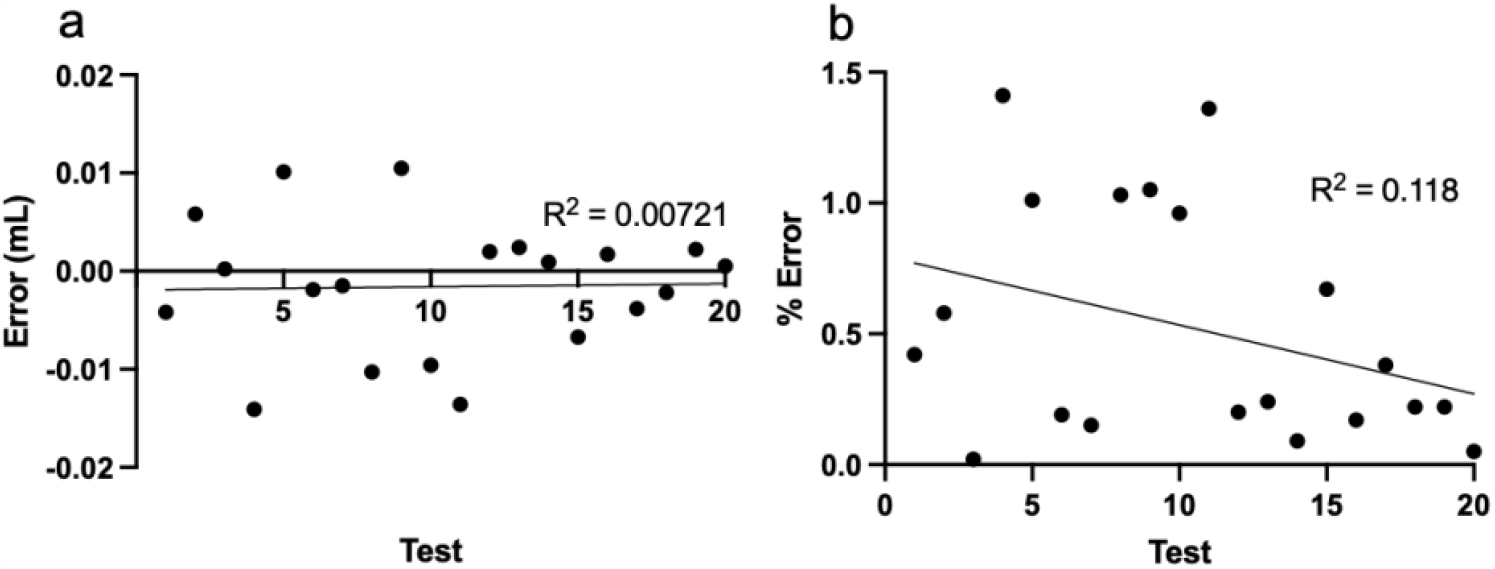
Characterization of pump error in **a)** mL, and **b)** % error. Neither showed significant non-zero slope indicating consistent error across tests. Average pump error was 0.5% (95% CI 0.3-0.7)

### Individual Fluid Consumption

Individual fluid intake binned by hour and averaged across all nine days is shown in figure 4. All rats showed greater ethanol consumption compared to water (Rat 1: F(1,16)=90.81, P<0.001; Rat 2: F(1,16)=137.9, P<0.001; Rat 3: F(1,16)=161.7, P<0.001; Rat 4: F(1,16)=4.817, P=0.043) and fluid consumption increased with time (Rat 1: F(6.161,98.57)=8.151, P<0.001, Rat 2: F(5.713,91.41)=6.535, P<0.001; Rat 3: F(6.324,101.2)=2.747, P=0.015; Rat 4: F(23, 368)=5.620, P<0.001). In 3 of the 4 rats, there was also an interaction effect (Rat 1: F(23,368)=2.942, P<0.001; Rat 2: F(23,368)=3.056, P<0.001; Rat 3: F(23,368)=1.857, P=0.01), with alcohol consumption increasing more than water over time.

**Figure 4.**
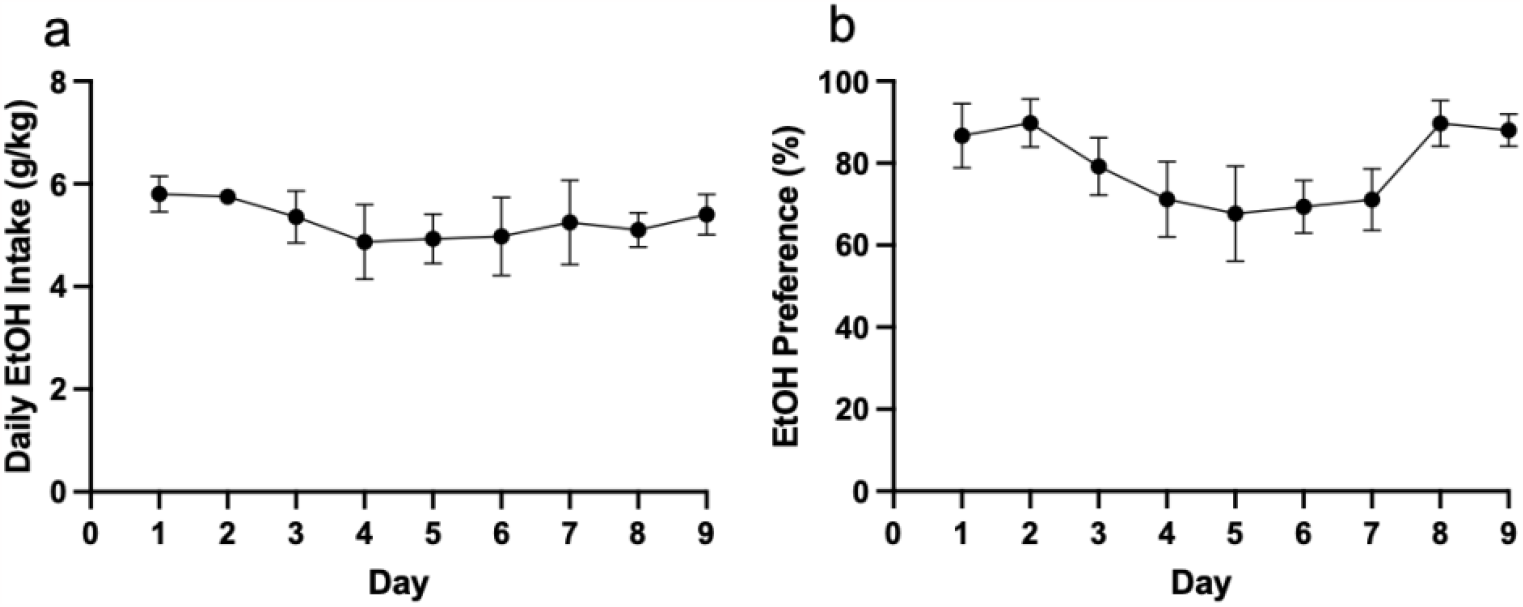
Average alcohol (10% ethanol v/v) consumption in group housed rats shown as **a)** daily intake (g/kg), and b) preference (%). Daily intake was consistent across days. Preference showed a significant effect of day, indicating shifting preferences across days, though no difference survived post-hoc.

### Circadian Effects on Fluid Consumption

Rats consumed a much greater volume of alcohol during the dark cycle (F(1,64)=494.7, P<0.001) as shown in figure 5a. Rats also consumed more water during the dark cycle (F(1,64)=45.74), P<0.001) as shown in figure 5b. Rats showed robust individual differences in fluid consumption of both alcohol (F(3,64)=7.397, P<0.001) and water (F(3,64)=17.78, P<0.001).

**Figure 5.**
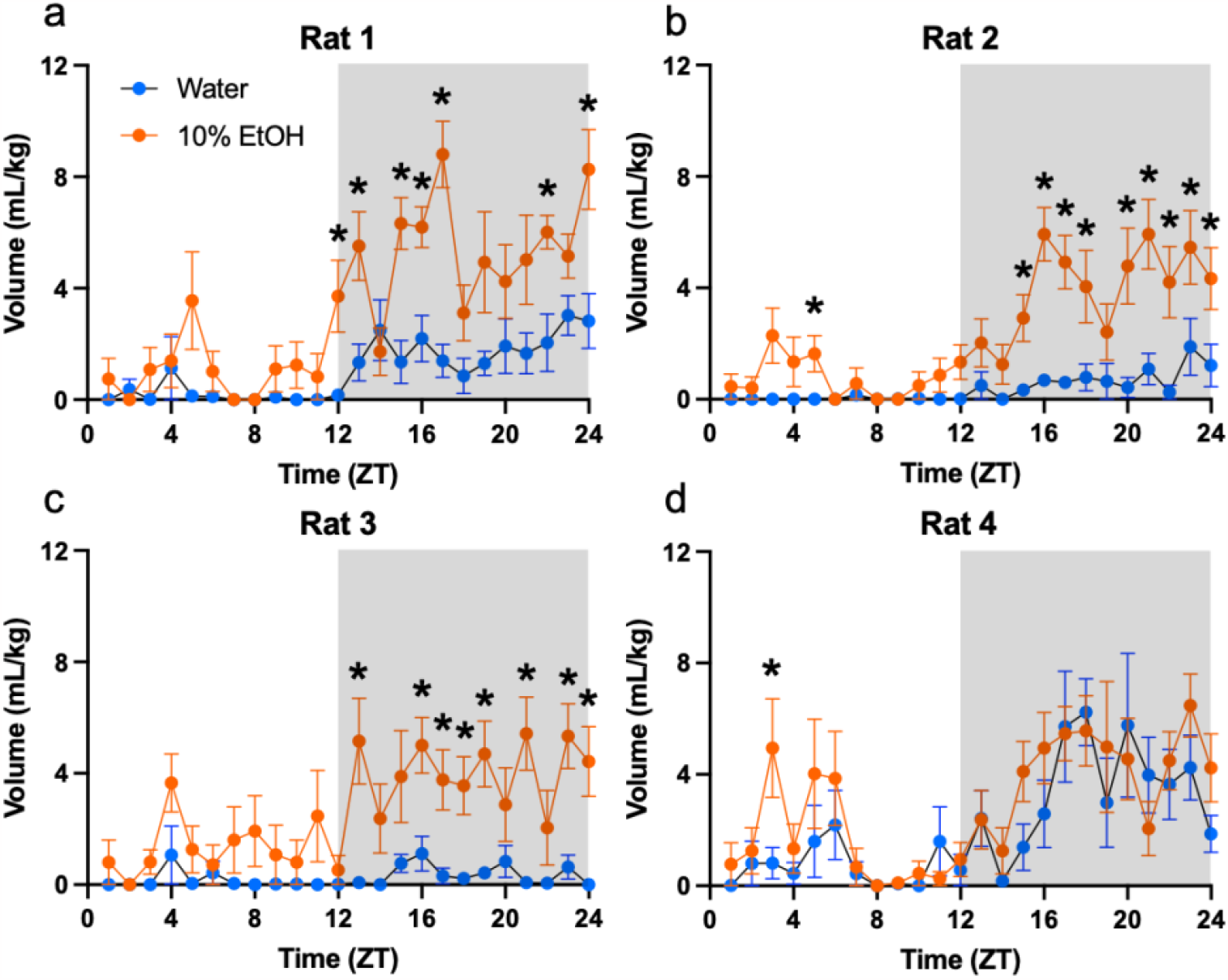
Individual fluid consumption binned by hour and plotted by Zeitgeber time with grey shading indicating the dark cycle. All rats drank more over time and consumed more alcohol than water. Rats 1-3 (a-c) also had a time by fluid-type interaction. *p<0.05 versus water.

### Fluid Type Effects on Bout Microstructure

Drinking was greatly affected by the fluid type, with the alcohol resulting in greater average bout size (figure 6a; t(3)=3.659, P=0.0353), max bout size (figure 6b; t(3)=7.088, P=0.0058), and volume per lick (figure 6c; t(3)=5.255, P=0.0134). Bout frequency also appeared to be higher but did not reach significance (figure 6d; t(3)=2.455, P=0.0913). The alcohol volume contributed by different bout ranges were also consistently greater than water (figure 6e; effect of fluid type: F(1,48)=37.83, P<0.001). Of most interest, there was an interaction between the bout range and the normalized volume consumed of each fluid type (figure 6f; F(7,48)=7.645, P<0.001), with greater normalized water consumption in small bouts, and greater normalized alcohol consumption in large bouts.

**Figure 6.**
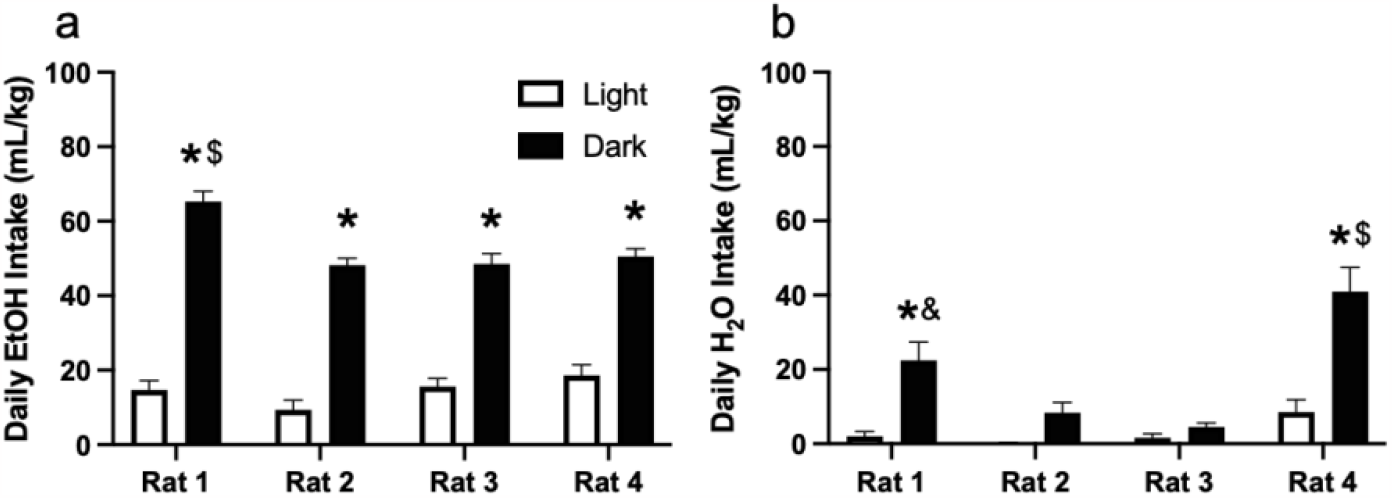
Average daily **a)** alcohol and **b)** water consumption during the light and dark cycle. Rats consistently drank more during the night cycle. *p<0.05 dark versus light. $p<0.05 versus all groups. &p<0.05 vs rat 2 and 3 dark.

**Figure 7.**
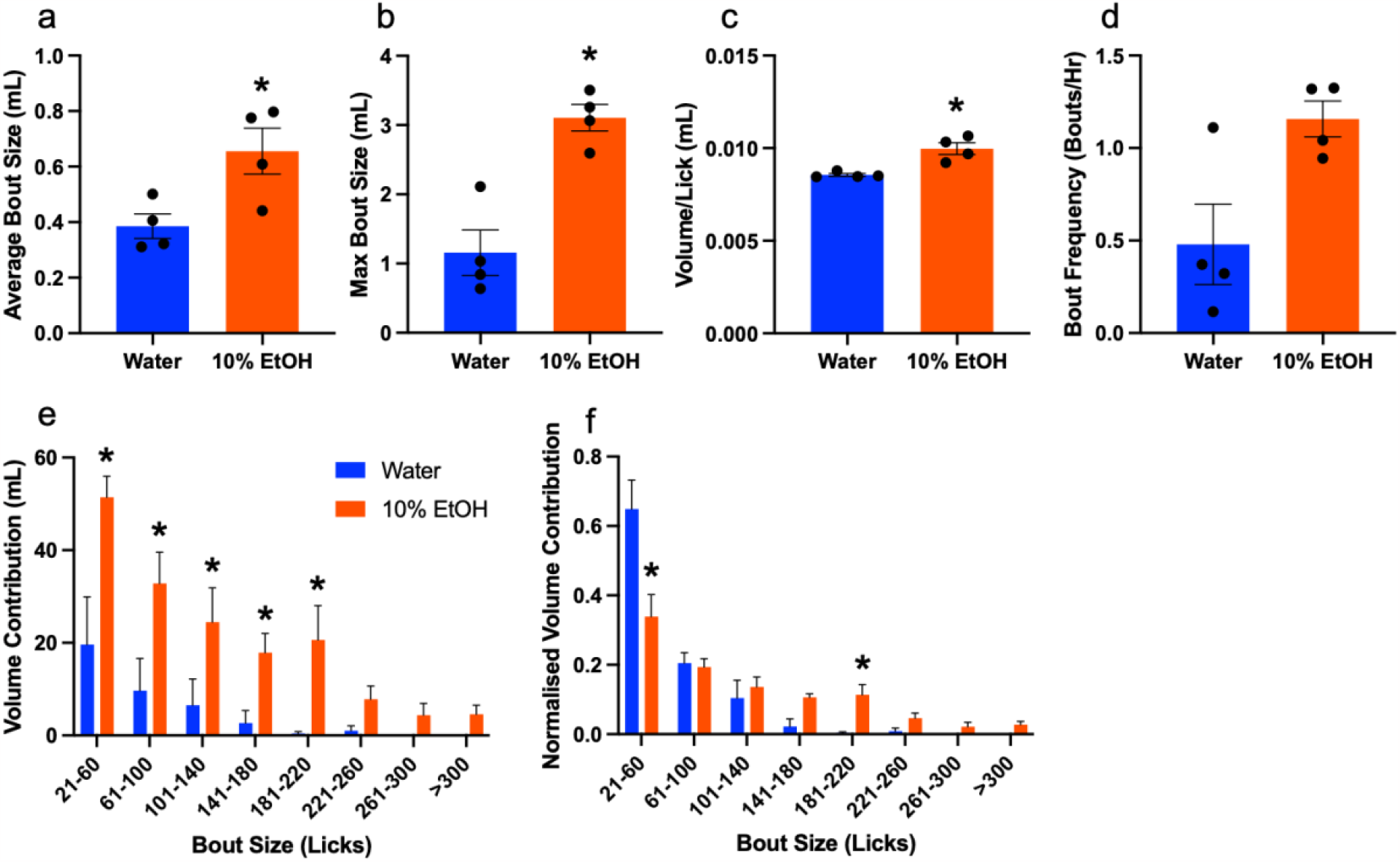
Bout structure analysis based on fluid type. Alcohol drinking resulted in greater **a)** average bout size, **b)** max bout size, and **c)** volume per lick. **d)** Bout frequency is marginally higher in alcohol compared to water (t(3)=2.455, P=0.0913) **e)** Overall volume consumed binned by lick number. A greater volume of alcohol was consumed across bout rangess. **f)** Volume contributed by each bout range normalized by the total volume of each fluid type. Normalized water consumption is higher than alcohol in small bouts, while normalized alcohol consumption is higher in larger bouts. *p<0.05 alcohol versus water.

## Discussion

Accurate tracking of fluid consumption is important in preclinical studies of substance use, as well as psychopathology (e.g., sucrose preference test). The majority of prior studies have individually housed animals to measure a subject’s fluid intake, thereby modeling consumption in a potentially-stressful, non-naturalistic, socially isolated environment. Therefore, to study fluid consumption disambiguated from isolation, social tracking of fluid consumption is required, especially when wanting to understand social determinants of drinking behaviour. Here, we developed a fully open-source implementation of a robust solution to social fluid consumption tracking that was able to show individual differences in alcohol preference, circadian influence on drinking, and fluid type-dependent bout structure.

Firstly, while there have been prior iterations of open-source peristatic pumps, our design presented here provides an alternative with remarkably accuracy. Using this device allowed us to track the volume consumed, rather than use a proxy for consumption such as licking. However, licking was still an important parameter, as we discovered a rat’s presence or beam-breaking was not adequate to define a drinking bout. Activation of the pump needed to correspond to licking to avoid false activation of the pump when a rat was near the RFID reader. We therefore used a capacitive lickometer on a chip design. Though infrared beam breaks have been used successfully as lickometers previously (Frie and Khokhar, 2019; Godynyuk et al., 2019), Petersen et al. have shown the superiority of capacitive systems over infrared as well as the improved flexibility of capacitive sensors on a chip compared to floor plate-based designs (Petersen et al., 2023). Thus, we were able to begin fluid delivery at a bout’s true start by activating the pump when a rat was both present and licking.

Pairing the lickometer with an Arduino-based microcontroller also allowed for time as an added dimension for fluid tracking. This enabled bout and circadian analysis of drinking behaviour. Rats were observed to do nearly all fluid consumption during the dark cycle as has been seen previously (Zucker, 1971). We were also able to show that rats tended to take larger bouts when drinking alcohol compared to water. Bout size may be an important measure related to excessive intake, with larger bout sizes in non-human primates being predictive of future heavy drinking (Grant et al., 2008), and greater bout sizes in high drinking rat lines (Samson, 2000). Quantity-frequency measures of alcohol consumption is also used in human diagnostics of regulatory loss of control (Feunekes et al., 1999).

The primary strength of using an RFID-based system is to track individual drinking bouts. FARESHARE was able to detect individual differences in drinking behaviour that would otherwise be lost if a simple group average was taken. One rat was observed to consume significantly more alcohol than its cage mates while another showed nearly no preference for alcohol over water, demonstrating FARESHARE’s ability to elucidate potential effects of rank, pretreatment, or individual substance vulnerability in a social context. As access is controllable via RFID, certain rats can have access restricted, allowing for control of partner rat consumption. Additionally, since the lickometer can still be active when access is restricted, extinction burst-like lick-spout activity measures may be taken when licking occurs without fluid delivery. Cage crowding can also be evaluated, as the system can handle as many animals as desired. FARESHARE is designed to be used in nearly any existing caging systems, therefore lacking the need to modify cages. As fluid reservoirs are outside the cage, frequent refilling can be avoided, preventing unwanted fluid loss, which is already reduced by active fluid delivery.

FARESHARE is not without limitations. The effect of viscosity on pump accuracy has not been evaluated. It is likely that highly viscous fluids will not work properly due to the low diameter of the tubing used, leading to inaccurate volume measurements or pump failure. Additionally, at the time of this experiment, the software did not allow for the measurement of some common microstructure measurements such as interlick interval, interbout interval, bout duration, or the direct measure of lick frequency, however, this has now been updated and implemented. Though the pump durability is quite good for a non-commercial product, if FARESHARE is to be used for extended periods such as water tracking over several months, the tubing will likely require replacement every couple of months to ensure there is no failure.

As FARESHARE is open-source, we are excited to see how labs alter the design to fit their specific needs and overcome any limitations that they may encounter. We also hope that the low-cost will allow more labs to implement social tracking of fluid intake and study social determinants of substance use, as these important factors are often ignored in traditional methods and provide translational validity. In conclusion, accurate tracking of social fluid consumption in preclinical studies of substance use is crucial for understanding social determinants of drinking behaviour. Prior studies have often individually housed animals, potentially inducing stress and social isolation, which can confound the results and overlooks a major translationally relevant aspect of substance use development and treatment. To address this issue, a fully open-source implementation called FARESHARE was developed, enabling social fluid consumption tracking, and allowing researchers to explore individual differences in alcohol preference, circadian influence on drinking, and fluid type-dependent bout structure. FARESHARE has the potential to become an invaluable tool in unraveling the complex dynamics of substance use and psychopathology, offering new insights into the social determinants of drinking behaviour and facilitating the development of more effective interventions.

